# IL-27 Stablizes Myc-Mediated Transcription In Memory-Fated, Vaccine-Elicited CD8+ T Cells

**DOI:** 10.1101/2024.07.31.606026

**Authors:** Scott B. Thompson, Michael Harbell, John Manalastas, Daria L. Ivanova, Kent A Riemondy, Erika Lasda, Vincent Chen, Jay R. Hesselberth, Anthony Phan, David Christian, Christopher A. Hunter, Tonya Brunetti, Laurent Gapin, Jared Klarquist, Ross M. Kedl

## Abstract

Protection from pathogens relies on both humoral (antibody-mediated) and cellular (T cell-mediated) responses. While infections robustly elicit both types of immunity, currently approved vaccine adjuvants largely fail to induce T cell responses on par with that instigated by infections. Our goal was to investigate the transcriptional programming that supports the formation of CD8^+^ T cells elicited by subunit vaccines compared to those elicited by infections. Our data show that vaccine-elicited T cells represent a transcriptionally unique subset of activated T cells with high proliferative capacity and a memory cell fate. This relies on IL-27 signaling, which stabilizes c-Myc and thereby supports the biomass acquisition necessary for clonal expansion. Collectively, our findings reveal that subunit vaccine-elicited T cells uniquely combine aspects of both memory and effector T cell subsets, and selectively utilize IL-27 signaling to sustain the clonal expansion of cells dedicated to a memory fate.

**One Sentence Summary:** In contrast to infection, subunit vaccines induce a distinct population of CD8+ T cells with memory fate characteristics which maintain their proliferative capacity during the expansion phase through IL-27 receptor signaling.

## INTRODUCTION

The overwhelming success of mRNA vaccines during the SARS-CoV2 pandemic have placed vaccines into the public consciousness comparable to the days of smallpox eradication and the polio epidemic. Despite their success in eliciting antibody, these vaccines, like all other approved subunit vaccine platforms, induce limited expansion of T cells even after multiple rounds of boosting ^1,2^. While this is largely acceptable for eliciting prophylactic humoral immunity, successful therapeutic intervention against chronic infectious diseases and cancer will inevitably require the elicitation of T cell responses that are orders of magnitude higher ^3^. A better mechanistic understanding of adjuvant-elicited immunity is necessary in order to successfully address the unmet clinical need of therapeutic cell-mediated immunity.

Our mechanistic understanding of robust cellular immunity has largely come from the study of immune responses to infectious challenge. It is generally believed that these mechanistic insights have direct application to the design, development, and formulation of subunit vaccines. However, there are striking differences in the importance of various cytokines (IL-27, IL-15), transcription factors (Tbet, FOXO1, Eomes, TCF1), and metabolic programs (oxidative phosphorylation) in vaccine-elicited versus infection-elicited CD8+ T cells responses ^4–7^. This suggests that mechanistic insights gained from infectious models cannot be assumed to apply directly to vaccine adjuvant-elicited cellular immunity.

c-Myc (Myc) is a well described transcription factor, with important roles in cellular proliferation, survival, metabolic function, and oncogenic transformation^8–11^. Myc has canonical functions as a broad amplifier of transcription and translation as well as a mediator of survival by facilitating aerobic glycolysis^8,9,12^. Transcription of Myc mRNA is temporary in activated T cells after antigen encounter ^13,14^, requiring post-transcriptional regulation for sustained activity. Under the appropriate conditions, Myc resists proteasomal degradation and sustains the expression of biomass-acquiring nutrient receptors, biosynthesis of mitochondria, and maintenance of mitochondrial health through mitochondrial fusion^8,10,11,15–18^. This tight regulation of Myc activity has three major advantages. First, it ensures that T cell proliferation is sustained only in an inflammatory environment. Under conditions of auto-antigen encounter without inflammation, the loss of Myc and its transcriptional targets results in a rapid cessation of clonal expansion, facilitating tolerance instead of immunity. Second, the same inflammatory cues that sustain Myc post-translationally^13,14^ also direct T cell differentiation toward effector phenotype and function and away from memory cell fates. Consequently, clonal expansion occurring more than three days after antigen encounter arises almost entirely from the effector, not memory, T cell pool. Third, the post-translational regulation of Myc function ensures that clonal expansion stops in conjunction with the cessation of inflammation. As Myc-sustaining inflammatory cues dissipate during infection resolution, effector T cells cease to divide and eventually die. In contrast, memory-fated T cells which opted out of rapid cell division earlier engage alternate transcriptional programs to support their longevity ^7,19^.

This mechanism of controlling Myc-mediated transcription in T cells has likely been evolutionarily selected by an infectious process in which inflammation begins subtly, grows steadily, and may take days or weeks to resolve. Its utility is far less clear after adjuvanted subunit vaccination, where both antigen and inflammation peak early and sharply dissipate within hours ^7^. In the present study, we explored whether T cell expansion in adjuvant and infectious contexts can be differentiated at the level of transcription. Building on our previous work, we used single cell RNA sequencing (scRNA-seq) to identify a unique transcriptional signature at four days post-vaccination, exhibiting features of clonal expansion and memory cells. The functional consequences of this transcriptional signature were confirmed *in vivo*, where rapidly proliferating T cells were predominantly memory-fated after subunit vaccination but nearly all effector-fated after infection. Finally, we identified IL-27 signaling as uniquely critical after subunit vaccination for maintaining Myc transcriptional activity, a prerequisite for biomass acquisition and subsequent clonal expansion.

## RESULTS

### Heterogeneity between the adjuvant- and infection-elicited responses is exhibited early after vaccination/challenge

We examined the response of OT1 CD8+ T cells to a polyIC/αCD40 combined-adjuvant (Adj) subunit vaccine, infection with *Listeria Monocytogenes* (LM), or infection with vaccinia virus (VV) by single-cell RNA-sequencing (scRNA-seq) at 4, 7 and 100 days after challenge (**Fig 1A**). Adoptively transferred OT1 T cells responding to ovalbumin (ova) were used to eliminate any potential confounding effects of differential TCR usage. A total of 3677 cells (Adj-1217, LM-1450, VV-1010) passed quality control (see Materials and Methods) and were integrated into a combined reference dataset. To identify and characterize subpopulation structures, we used unsupervised graph-based clustering, with distinct niches separated by time and challenge. (**Fig 1B-D**). These distinctions were generally lost by day 100, where cells of different origins were homogeneously mixed (**Fig 1D**). Using a shared-nearest-neighbor approach to identify clusters of cells with closely related transcriptional profiles, sixteen gene expression clusters were characterized (**Fig 1E-F**). As anticipated, all clusters associated with the day 100 time point (clusters 11-16) showed nearly equal distribution of Adj, LM and VV-elicited cells (**Fig 1F**). Consistent with their presence at a memory time point, these clusters showed high expression of genes associated with T cell memory formation (*Il7r*, *Dapl1*, *Tcf7*), and low expression of effector-associated transcripts (*Zeb2*, *GrzB*, *Klrg1*) (**Fig 1G-H and Fig S1**). Examining earlier timepoints revealed greater degrees of heterogeneity between modes of antigenic challenge, where some clusters were predominantly comprised of cells derived from a single challenge (**Fig 1F-G**). The adjuvanted vaccine-elicited (Tvac) cells in cluster 2 were particularly intriguing, as they had elevated expression of memory fate-associated genes (most notably *Tcf7*) within 4 days of vaccination (**Fig 1H-I**). In contrast, infection-elicited T cells (Tinf) from LM or VV challenge showed early skewing toward effector-fated gene transcription (**Fig 1H-I**). These extend our previous analysis of adjuvant elicited T cells at 7 days post vaccination ^7^ and suggest that the majority of Tvacs are pre-disposed toward a memory fate even earlier than previously expected.

**Fig 1.**
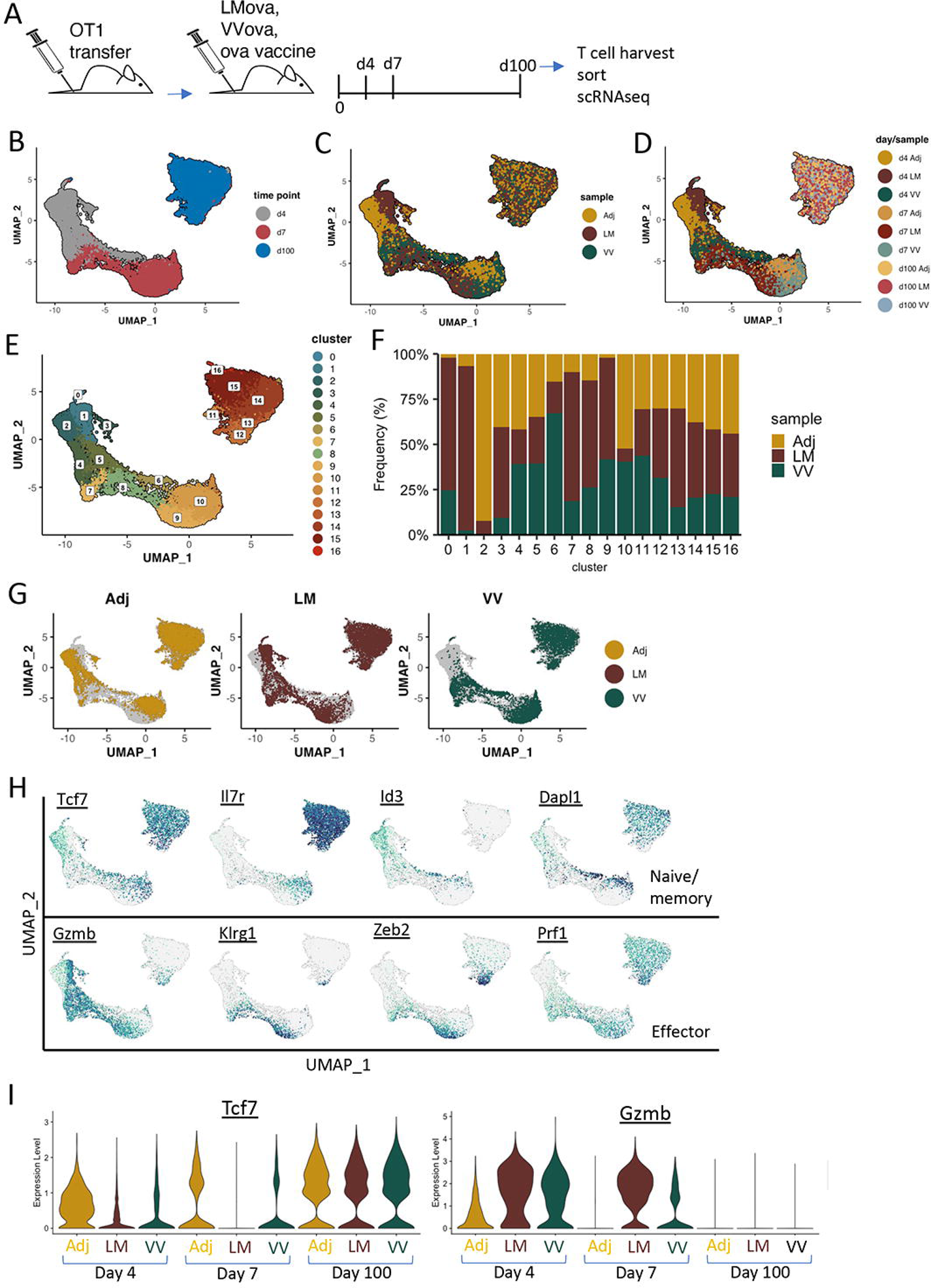
Heterogeneity between the adjuvant- and infection-elicited responses is exhibited early after vaccination/challenge. A. Schematic of experimental design to capture scRNA sequence from CD8+ T cells isolated at days 4, 7 and 100 post vaccination/challenge from vaccine, VV or LM. B-E. UMAP of samples derived from all vaccination/challenge time points, highlighted by A. time point, B, immunological challenge/sample, C. combined day/sample, and D. UMAP clustering. F. Fraction of each UMAP cluster represented by each sample. G. UMAP split by immunological challenge H. UMAP highlighted for genes associated with memory (top) or effector (bottom) fate designation. I. Violin plots highlighting the relative expression level of TCF7 (left) or Gzmb (right) for each sample and at each time point.

### Tvacs display an early dual commitment to proliferation and a memory cell fate

Reclustering of day 4 scRNA-seq data confirmed distinct transcriptional states for Tinfs and Tvacs at this early time point. Clusters 1 and 3 were mostly T cells derived from LM or VV challenge, respectively, whereas cluster 2 was again comprised almost entirely of Tvacs (**Fig 2A-C**). Cells were largely undergoing cell cycle progression (S/G2M) regardless of challenge (**Fig 2D**), though the phenotype of proliferating Tvacs differed significantly from cells derived from either infectious challenge. As expected, Tinf in S/G2M expressed markers consistent with an effector cell fate (exemplified by *Gzmb),* while expression of markers of memory-fate (such as *Tcf7*) were only expressed for cells in G1 (**Fig 2E**). In contrast, Tvacs expressed high levels of *Tcf7* regardless of their cell cycle stage (**Fig 2E**). This indicates that while memory-fated Tinf cells exit the cell cycle, Tvacs are uniquely able to sustain clonal expansion while maintaining a memory cell fate in the earliest rounds of cell division.

**Figure 2.**
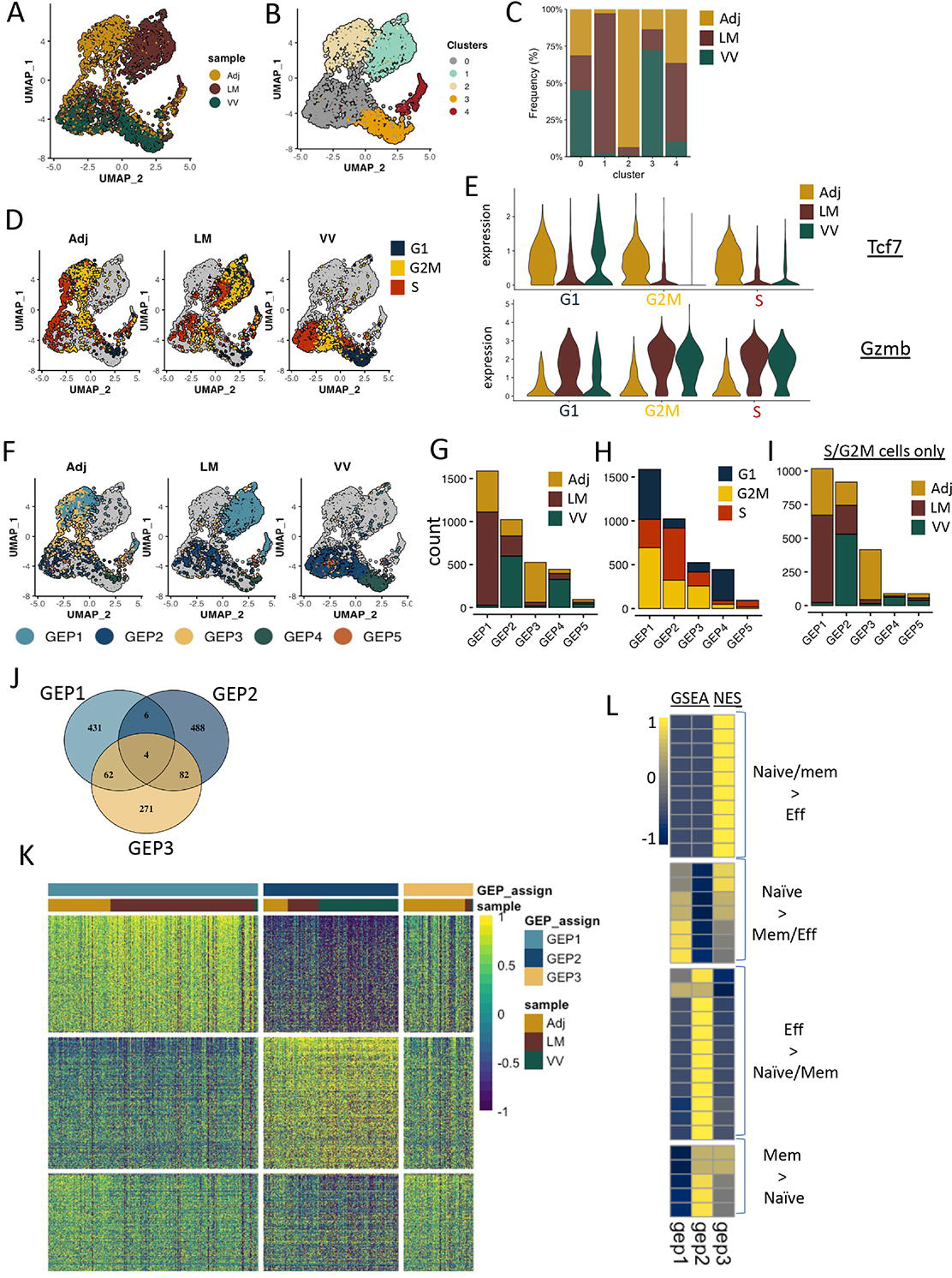
Tvacs display an early dual commitment to proliferation and a memory cell fate. A, B. UMAP for only day 4 scRNA analysis, highlighted for A) immunological challenge/sample, and B) UMAP clustering. C. Representation of cells derived from each immunological challenge/sample within each cluster. D, E. UMAP highlighted by cell cycle phase, split by D) cell cycle phase, or E) immunological challenge/sample. F. Violin plots for average expression of Tcf7 (top) or Gzmb (bottom) for cells in each cell cycle phase split by immunological challenge/sample. G. UMAP highlighted by cNMF assigned gene expression programs (GEPs). H. Representation of cells derived from each immunological challenge/sample within each GEP. I. Representation in each GEP for cells in each cell cycle phase. J. Venn representation (top) and percentage overlap (bottom) in expression of GEP-defining genes between cells assigned to GEPs 1-3. K. Heatmap for expression of all GEP-defining genes from GEPs 1-3. Top color bar identifies the assigned GEP and sample designation of each cell. L. Gene set enrichment analysis (GSEA) of GEP-defining genes (for GEPs 1-3) against gene sets specific to CD8+ T cell activation/differentiation (see Table S1 for specific gene sets)

Although scRNA-seq quantifies transcripts in individual cells, each cell’s expression profile can be a composite of both cell-type identity and cellular activity, complicating their separation. We employed consensus non-negative matrix factorization^20^ on the day 4 samples to infer identity and activity programs, identifying five gene expression programs (GEPs) (**Fig. 2F**). Similar to clustering, these five GEPs predominantly consisted of cells from LM (GEP1), VV (GEP2 and GEP4), or Adj (GEP3) (**Fig. 2F-G**). Analyzing the cell cycle phases of cells within each GEP revealed that the majority of cells in the S/G2M phase were confined to GEPs 1-3 (**Fig. 2H**). This observation was further confirmed by evaluating only the actively cycling cells (**Fig. 2I**). Given our primary interest in the interplay between cell fate and proliferation, we focused our subsequent analyses on GEPs 1-3.

To determine GEP-defining genes, we plotted gene ranking against the gene_spectra_score output from the cNMF analysis. These rankings were fitted to a sigmoid curve to calculate the minimum threshold for genes to be retained in a given GEP (**Fig. S2 and Table S1**). Upon comparing the gene sets, it was apparent that while the genes associated with GEPs 1 and 2 were unique relative to one another, they both had a noticeably increased overlap with GEP3 (**Fig 2J**). This was confirmed by evaluating the expression of the individual genes that compose each GEP, where the distinction between cells expressing of GEP1- and GEP2-defining genes is as notable as the similarity of GEP3-expressing cells for the genes expressed in GEP1 and GEP2 (**Fig 1K**). Next, we performed Gene Set Enrichment Analysis (GSEA) for the GEP-defining genes against the mouse C7 Immunologic gene signature gene sets. From all CD8+ T cell associated gene sets identified as statistically significant (**Fig. S3**), specific patterns were associated with each GEP. GEP2 genes were enriched in gene sets associated with effector cells as compared to either naïve or memory (**Fig 2L and S3**), consistent with more terminally-differentiated cells. GEP1 genes were represented in some effector gene sets, but were more associated with less committed, naive T cell gene expression (**Fig 1L and S3**). Lastly, GEP3 gene sets not only overlapped with a number of GEP1 and GEP2 gene sets, but also overlapped with numerous gene sets over-represented in naive/memory versus effector T cells.

In addition to GSEA, we examined the association of each set of GEP-associated genes with Gene Ontology-Biological Processes (GO-BP) pathways. As anticipated, most of the statistically significant pathways were related to the cellular and metabolic demands of cell division (**Fig. S4A and Table S3**). GEPs 1 and 2 appeared to be executing opposite biological processes, such that a process present in GEP1 was either absent or negatively associated with GEP2 (**Fig. S4A**). In contrast, GEP3-associated biological pathways showed substantial overlap with those of both GEPs 1 and 2 (**Fig. S4A**).

Finally, we used single-cell regulatory network inference and clustering (SCENIC) to predict transcription factor activity for each set of GEP-defining genes. Whether evaluating all transcription factor (TF) regulons (**Fig. S4B and Table S4**) or a representative set (**Fig. S4C**), the previous themes were reaffirmed; GEP1 and GEP2 predominantly expressed opposing TF regulons, while both showed considerable overlap with GEP3 TF regulons.

Since the GEP3 consists almost entirely of Tvacs, the GSEA, GO-BP and SCENIC analysis collectively reinforce the conclusion that adjuvant-elicited T cells express transcriptional programs commitment to a memory cell fate and rapid proliferation/clonal expansion. Further, these data indicate that Tvacs adopt this dual commitment well before the peak of the response ^7^. In contrast, clonally expanding Tinfs exhiibit either the conventional and expected hallmarks of short-lived effector T cells (VV, GEP2), or early effector T cells yet to diverge into separate memory or effector cell fates (LM, GEP1).

### Tvacs sustain proliferation of TCF1^hi^ cells, Tinfs sustain proliferation of TCF1^lo^

We further examined the consequences of these transcriptional analyses in vivo using a combination of EDU incorporation and cell proliferation dye labelling. To evaluate the phenotype at the earliest rounds of cell division, OT1 T cells were dye labelled and transferred into congenically distinct B6 recipients, which were then vaccinated or infected with LM. Harvesting mice at different time points over 72 hours reliably identified cells in divisions 0-9. We defined a region for cells expressing high levels of TCF1 across all cell divisions (**Fig 3A**), then determined the ratio of TCF1^hi^/TCF1^lo^ cells within each division (**Fig 3B**). After LM infection, equal numbers of T cells expressed high and low TCF1 up through division 4-5. Subsequently, the TCF1^hi^/TCF1^lo^ ratio rapidly dropped to less than 0.1 (**Fig 3B**), indicating that only the TCF1^lo^ cells continued to progress through cell divisions. In contrast, TCF1^hi^ Tvacs outnumbered TCF1^lo^ Tvacs ∼10:1 from divisions 1-7. Thereafter, TCF1^hi^ cells continued to dominate the response until the very last cell division identifiable where they matched the frequency of more effector-fated TCF1^lo^ cells.

**Figure 3.**
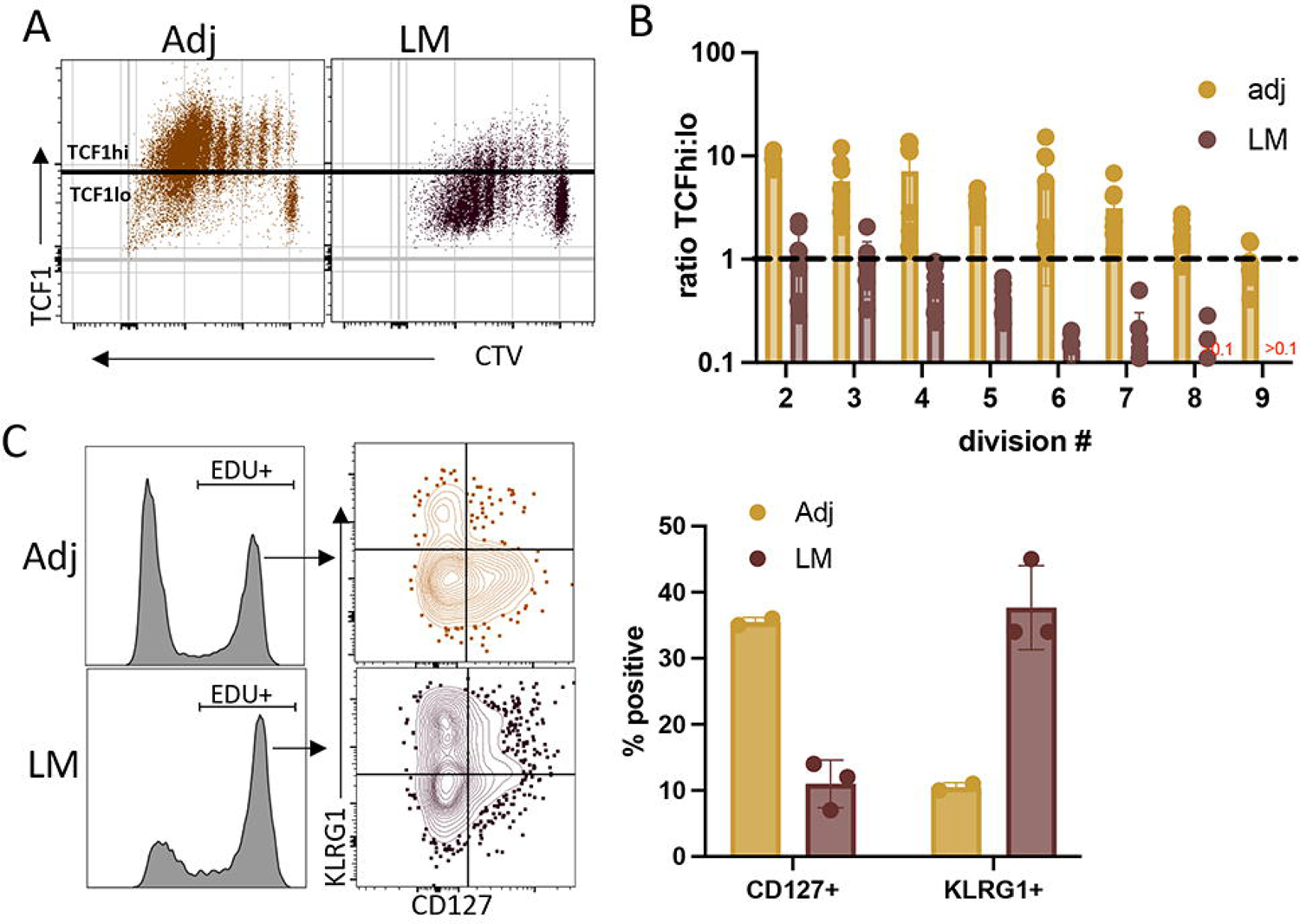
Tvacs sustain proliferation of TCF1^hi^ cells, Tinfs sustain proliferation of TCF1^lo^. A, B. CD45.1/2+, CTV labelled OT1 T cells were transferred into CD45.2/2 B6 recipients which were then either immunized with ova protein and combined adjuvant (Adj) or challenged with LM-ova (LM). To capture divisions 2-9, spleens were harvested at 36, 48 and 60 hours post vaccination, or 60 and 72 hours post LM challenge. The expression of TCF1 (hi vs low) was evaluated at each cell division. A. Dot plot of concatenated files from each time point showing the expression of TCF1 over divisions 0-9. B. Quantification of A per division, each dot representing individual mice with >20 cells in each division.. C, D. B. WT CD45.1/2+ OT1 T cells were transferred into CD45.2/2 B6 recipients which were then either immunized with ova protein and combined adjuvant (Adj) or challenged with LM-ova (LM). Five days later, the mice were injected IP with 200ug EDU and 2 hours later sacrificed and the transferred T cells in the spleen were evaluated for the CD127 x KLRG1 phenotype of all T cells that had incorporated EDU. C. Representative histograms of EDU incorporation and KLRG1 x CD127 staining for EDU+ OT1 T cells. D. Quantification of the frequency of cells in the lower right (CD127+) and upper left (KLRG1+) quadrants. Data are representative of three experiments performed. All data shown are mean ± SEM; n= 3-5 mice per group, representative of 3 independent experiments. Significance was defined by unpaired t test with Welch’s correction and two-way ANOVA, where *p < 0.05, **p < 0.01, ***p < 0.001.

The enhanced proliferative capacity of memory-fated Tvacs was additionally identified by EDU incorporation at later time points when CTV labeling was no longer effective. Five days after vaccination or LM-ova challenge, mice were injected with EDU, which incorporates into DNA during replication, two hours prior to sacrifice. Following cell surface staining, EDU+ cells were identified by flow cytometry, and the EDU+ versus EDU− cells were evaluated for their expression of memory versus effector cell fate markers. EDU incorporation was predominantly observed in the CD127lo KLRG1hi short-lived effector cells (SLECs) after LM infection, whereas a substantial portion of EDU+ Tvacs bore the memory precursor effector cells (MPECs) phenotype of CD127hi KLRG1lo (**Fig. 3C**). Thus, the unique proliferative capacity of memory-fated Tvacs predicted by the ex vivo scRNA-seq analysis was confirmed in vivo.

### IL-27 signaling sustains GlcNacylation of Myc and its transcriptional activity in memory-fated Tvacs

IL-27 plays an obligate role in supporting the accumulation of CD4+ and CD8+ Tvacs but not Tinf ^4,5^, the mechanistic underpinnings of which have yet to be fully elucidated. A curious feature of this IL-27-dependency is its delayed onset; in a co-transfer setting, WT and L-27R^-/-^ OT1 T cells expand equally through day 3 post vaccination, after which the IL-27R^-/-^ T cell fail to accumulate further (**Fig 4A**) ^5^. Analysis of bulk RNA sequencing from WT and IL-27R^-/-^ T cells 3 days after vaccination revealed a dramatic loss of Myc transcriptional targets, readily apparent by GSEA (**Fig 4B**) or display of Myc target genes defined by the Hallmark gene signature (**Fig 4C**).

**Figure 4.**
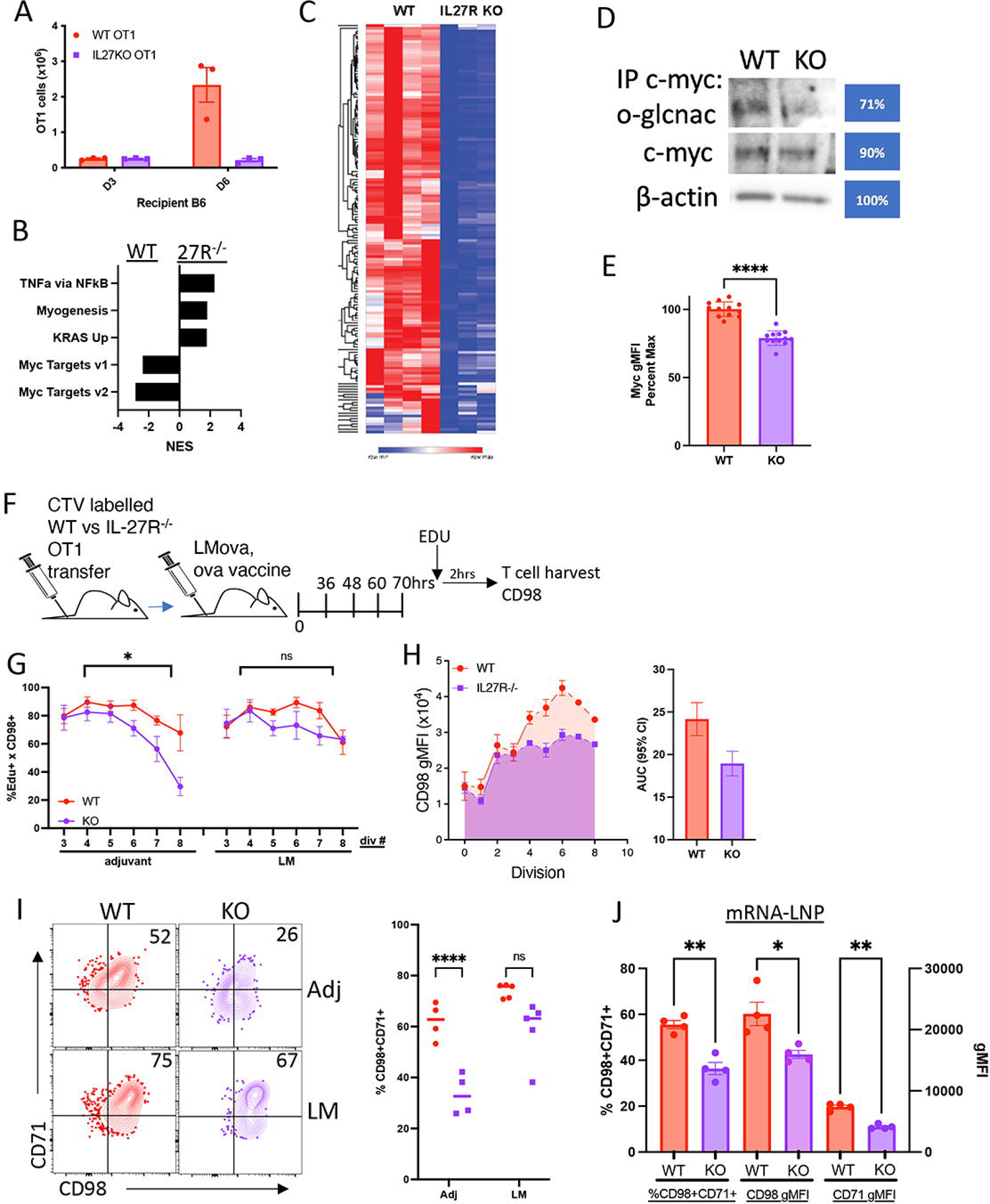
IL-27 signaling sustains GlcNacylation of Myc and its transcriptional activity in memory-fated Tvacs. A-I. WT CD45.1/2+, and IL-27R-/- CD45.1/1, OT1 T cells were co-transferred into CD45.2/2 B6 recipients which were then immunized with ova protein and combined adjuvant. A. Representative mice were sacrificed 3 and 6 days later and the number of OT1s in the spleen enumerated by flow cytometry. B. Gene Set Enrichment Analysis of bulk RNAseq from WT and IL-27R-/- OT1s flow sorted to purity at day 3 post vaccination. C. Heatmap of all Myc target genes defined by the Hallmark gene signature from WT and IL-27R-/- T cells. D. Western blot from flow sorted WT and IL-27R-/- OT1 T cells 3 days post vaccination. Top row: immunoprecipitation for cMyc, followed by immunoblot with GlcNacylation-specific antibody, RL2. middle row: western blot for cMyc of lysates from WT and IL-27R-/- OT1s. Bottom row: western blot for beta-actin of lysates from WT and IL-27R-/- OT1s. E. Flow cytomeytric analysis of Myc gMFI across 12 mice from 4 experiments, normalized by the mean Myc gMFI of WT cells in each experiment. F. Schematic of the experimental methods used in G-H, incorporating both CTV and EDU labelling of co-transfered, congenically distinct WT and IL-27R-/- OT1s. G. Spleens were harvested at 36, 48 and 60 hours post vaccination, or 60 and 72 hours post LM challenge. The frequency of EDU+ CD98+ cells was evaluated at each cell division. Statistical significance of genotype (p=0.0069, cell division (p=0.0004), and division x genotype (p=0.02) was determined by fitted mixed-effects analysis. H. Performed as in G, quantifying area under the curve (AUC) of the gMFI of CD98 at each cell division for cotransfered WT and IL-27R-/- T cells in vaccinated hosts. Error bars represent 95% confidence interval for AUC. Co-transferred WT and IL-27R-/- T cells from the spleens five days after vaccination or LM challenge. Dot plots show % of cells that are CD98+CD71+ (upper right quadrant) of all EDU+ cells, quantified in graph to the right. J. Experiment performed as in I, but spleen cells harvested 6 days post ova mRNA-LNP vaccination. Left-most bars and axis quantify frequency of CD71+CD98+ WT or IL-27R-/- T cells. Middle and right-most bars and axis quantify gMFI of CD98 or CD71 of WT or IL-27R-/- T cells. Data are representative of 2-4 experiments performed n= 3-5 mice per group. Significance was defined as stated, where *p < 0.05, **p < 0.01, ***p < 0.001, ****p < 0.0001.

Transcription of Myc RNA occurs largely in the first 72 hours post-TCR engagement ^13,14^, after which, the half-life of Myc protein can be stabilized through GlcNacylation^17^, a downstream consequence from activation of the Hexosamine Biosynthetic Pathway (HBP). Interestingly, direct interrogation of Myc four days after vaccination identified reduced GlcNacylation in IL-27R^-/-^ as compared to WT OT1 T cells (**Fig 4D**). This reduction correlated with an overall loss of Myc protein, evaluated by either western blot (**Fig 4D**) or flow cytometry (**Fig 4E**). Collectively, these data suggested that IL-27 might contribute to the activation of the HBP in Tvacs, resulting in the post-translational stabilization of Myc, thereby sustaining the transcription of genes central to biomass acquisition and metabolism.

To test this hypothesis, we evaluated the expression of the Myc target CD98 ^10,15^ on proliferation dye-labelled WT and IL-27R^-/-^ OT1 T cells responding to either adjuvanted vaccination or LM challenge (**Fig 4F**). Two hours before harvesting splenic T cells, the mice were injected with EDU to identify all cells actively replicating their DNA. In vaccinated hosts, IL-27R^-/-^ OT1 cells showed a significantly reduced proportion of EDU+ cells that were positive for CD98 after more than five cell divisions (**Fig 4G**), highlighting the necessity of IL-27 for sustaining the expression of Myc target genes in Tvacs. In contrast, and consistent with the IL-27-independency of the response to LM challenge ^4,5^, the frequency of EDU+ cells positive for CD98 in IL-27R^-/-^ T cells after infectious challenge was indistinguishable from that of WT T cells (**Fig 4G**). This difference in Myc activity was additionally notable when evaluating the area under the curve for CD98 gMFI x cell division (**Fig 4H**). The loss of Myc-dependent target expression in IL-27R^-/-^ OT1 T cells responding to vaccination continued to be apparent at later time points, particularly when incorporating CD71 expression as an additional proxy of Myc transcriptional activity ^10^. Whether evaluating the frequency of EDU+ cells positive for CD98+CD71+ (**Fig 4I**) or quantifying the expression of CD98 or CD71 on EDU+ T cells (**Fig S5A-B**), IL-27R^-/-^ Tvacs were substantially reduced relative to their WT counterparts. Again, there was no difference between WT and IL-27R^-/-^ T cells in the response to LM challenge.

We previously demonstrated that CD4+ and CD8+ T cell responses to a wide range of adjuvants are similarly dependent on IL-27 ^4^. This implies that IL-27’s role in maintaining Myc transcriptional activity is a fundamental mechanism underpinning the broader response to vaccination, not just the combined-adjuvant protein subunit vaccine used in these studies. Data presented by Phan et al. in this issue provide additional evidence that the cellular response to vaccination with mRNA-LNPs is also critically dependent on IL-27. Similar results have been obtained in our hands (**Fig. S5C**), leading us to investigate whether dysregulated expression of Myc-dependent targets is observed in IL-27R^-/-^ OT1s from mice immunized with mRNA-LNPs. Consistent with the results from the combined-adjuvant approach, EDU+ IL-27R^-/-^ T cells exhibited similar deficits in CD98 and CD71 co-expression following mRNA-LNP vaccination (**Fig. 4J**).

## CONCLUSION

Altogether, our results show that adjuvanted vaccines elicit T cell responses with a unique transcriptional program, coupling T cell proliferation with a memory cell fate. This is in striking contrast with the T cell responses to infections in which clonal expansion and memory fate designation are separated. In such setting, proliferating cells retain an effector fate while memory-fated cells exit the cell cycle. This unique Tvac program requires IL-27R signaling, which we can now mechanistically link to the capacity for IL-27 to post-translationally stabilize Myc activity, leading to the downstream expression of genes necessary for biomass acquisition and cellular division.

Linking IL-27 signaling to the sustained regulation of Myc provides some answers to long-standing questions about the control of clonal expansion in T cell responses. More than 20 years ago it was established that CD8+ T cells respond to initial antigenic stimulation by instigating a pre-established set of cellular divisions, even when the initial TCR stimulus is rapidly withdrawn ^21–24^. However, the longer T cells are allowed to interact with antigen, the larger their peak expansion ^21,25^. If T cells are pre-programed to respond to a TCR stimulus regardless of its duration, how can more sustained TCR engagement be relevant to the peak response? These seemingly contradictory observations can coexist because of the transcriptional and post-translational regulation of Myc ^10,13–15^. Transcription of Myc RNA is induced following TCR stimulation, and while both the strength of the TCR stimulus and the nature of the costimulatory and cytokine environment can influence the magnitude of mRNA that is transcribed ^13,15^, the duration of transcription is limited to ∼72 hours ^15^. By virtue of the genes that it targets for expression, Myc is absolutely necessary for biomass acquisition and blast formation ^10^. Unless post-translational modifications are made to Myc, GSK3B-mediated phosphorylation of threonine 58 results in Myc recruitment to, and degradation by, the proteosome ^26–28^. In the absence of any other signaling interventions, the relatively fixed window for transcription of Myc RNA transcription serves as a cell intrinsic “division timer”^13^, explaining how CD8+ T cells undergo a pre-determined number of cell divisions even after minimal TCR engagement.

To prolong clonal expansion, however, Myc protein must be stabilized post-translationally. This can be achieved through various mechanisms, one of the better characterized being the HBP. The network of enzymes mobilized by the HBP culminate in the activation of O-GlcNAc Transferase (OGT) and its GlcNacylation of threonine 58 on Myc, prohibiting the phosphorylation of the same residue by GSK3B and protecting Myc from proteasomal degradation ^17^. Multiple proinflammatory innate pathways and cytokines induce HBP activity ^15,17,29^, stabilizing Myc and prolonging clonal expansion through additional rounds of division. To this list of molecules that stabilize post-translational Myc protein through the HBP, we can now add IL-27, specifically in the context of adjuvanted vaccine-elicited cellular immunity.

The reasons for the specificity of IL-27 to adjuvanted vaccination for HBP activation and Myc stabilization remain unclear. However, this may be related to the unique proliferative demands of memory-fated T cells which most vaccine adjuvants produce in high frequency^7,30^. Ongoing work aims to integrate transcriptional and metabolic pathways into a more unified understanding of the immunological mechanisms unique to adjuvanted vaccine-elicited cellular immunity.

## RESOURCE AVAILABILITY

### Lead contact

Further information and requests for resources and reagents should be directed to and will be fulfilled by the lead contact, Ross Kedl.

### Materials availability

This study did not generate unique reagents. All materials used are available upon request.

### Data and code availability

Data reported in this paper will be shared by the lead contact upon request. This paper does not report original code. Any additional information required to reanalyze the data reported in this paper is available from the lead contact upon request. RNAseq and single-cell RNA-seq data have been deposited at GEO and are publicly available as of the date of publication. Accession numbers are listed in the key resources table. Flow cytometeric data are publicly available upon request as of the date of publication. DOIs are listed in the key resources table.

## EXPERIMENTAL MODEL AND SUBJECT DETAILS

### Mice

All experiments involving mice were conducted following protocols approved by the University of Colorado Institutional Animal Care and Use Committee (IACUC) according to guidelines provided by the Association for Assessment and Accreditation of Laboratory Animal Care. WT (C57BL/6J), congenic CD45.1 B6 (B6.SJL-Ptprca Pepcb/ BoyJ), ova-specific TCR-transgenic OT1 (C57BL/6-Tg(TcraTcrb)1100Mjb/J) mice were originally obtained from the Jackson Laboratory and subsequently bred inhouse at the University of Colorado Anschutz Medical Campus. IL-27Ra-/- mice were originally provided by Genentech ^31^. This study was carried out in accordance with protocols approved by The University of Colorado Anschutz Medical Campus Institutional Animal Care and Use Committee (IACUC), protocol #00172. Experiments were performed in 6 week-old or older male and female mice.

## METHOD DETAILS

### Immunization and infection

Mice were vaccinated (Adj) via tail vein injection (IV) with poly(I:C) (40 µg; InvivoGen) + αCD40 antibody (40 µg; clone FGK4.5, BioXcell), and detoxified whole chicken ova protein (150 µg; Sigma). This adjuvanted vaccine (Adj) was made immediately prior to immunization. For infectious challenge, mice were injected IV with 2 × 10^3^ colony-forming units (CFU) of *Listeria monocytogenes* expressing whole ova (LM), or 1×10^7^ PFUs of vaccinia virus expressing ovalbumin (VV).

### mRNA-LNP formulation

The full-length DNA sequence of ovalbumin with optimized 5’ and 3’ UTRs and in-frame 3X Flag-tag was synthesized by Twist Bioscience in a pTwist Kan High Copy plasmid vector. CleanCap AG capped mRNA, including N1-methylpseudouridine-5’-triphosphate substitutions, was produced by T7 RNA polymerase transcription from linearized plasmid templates and subsequently enzymatically polyadenylated with E. coli Poly(A) Polymerase. Column-purified mRNA transcripts were incorporated into LNPs composed of Precision Nanosystem’s Neuro9 lipid mix (catalog#: NWS0001) on a microfluidic LNP mixer (e.g., Precision NanoSystems Spark) and dialyzed twice with PBS (pH 7.4 without Calcium and Magnesium, Corning #: 21-040-CV). Encapsulation efficiency and RNA content of LNPs was quantified using the ThermoFisher RiboGreen RNA Assay Kit (catalog# R32700) a dye-binding assay following detergent-based dissolution of the LNPs and LNPs at concentrations of 50-100 ng/µl were used for in vivo administration. Ova-LNPs containing 1-2ug of mRNA were injected IV into recipient mice.

### Adoptive cell transfer

CD8+ T cells were magnetically purified from the spleens of indicated donor mice by negative selection using MojoSort Mouse CD8 T cell Isolation kit (Biolegend) to >95% purity. Where indicated, CD8+ T cells were CTV-labeled (Invitrogen) before transfer. OT1 cells were counted on a Vi-CELL automated cell counter (Beckman Counter) and indicated OT1 cell numbers (5 × 10^2^ or 5 × 10^3^) in 1 × PBS were transferred IV to recipient mice 1 day before or on the day of immunization or infection.

### Cell isolation

Where indicated, donor OT1 cells that were present at low numbers post adoptive transfer were magnetically enriched with magnetic columns (LS Columns, Miltenyi Biotec). For early (≤ 72 h after immunization or infection) time points, spleens were removed and crushed between glass microscope slides into 6 well plate containing 2 ml Click’s Medium (FUJIFILM Irvine Scientific) and 30 µl of 5 mg/ml Collagenase D (Roche) and 100 µl of 2 mg/ml DNase I (Worthington Biochemical) per spleen. After a 15-20 min incubation at 37°C, 2 ml 0.1 M EDTA in 1 × PBS was added and cells were incubated at 37 °C for additional 5 min. The cells were then washed with HBSS (Life Technologies) containing 5 mM EDTA and forced through 70 or 100 µm strainer. After wash, the cells were RBC lysed for 1 min in ACK buffer, then resusupended in complete RPMI and filtered through strainer. To enrich for CD45.1+ or CD45.2+ OT1, cells were resuspended in 500 µl RPMI and incubated with 10 ul of pulldown Ab (CD45.1 APC or CD45.2 APC) for 20 min at 4°C. After wash, cells were incubated in 500 µl RPMI with 5 ul anti-APC nanobeads (Biolegend) with rotation for 15 min at 4°C. After wash, cells were resuspended in 500 ul RPMI, filtered and purified through LS column (Miltenyi Biotec). For staining of spleenocytes at the later time points (> 72 h after immunization or infection), whole organs were crashed through 70 or 100 µm strainers into complete RPMI to generate single cell suspensions. The cell suspensions were then RBC lysed in ACK buffer, washed in complete RPMI and counted on a Vi-CELL (Beckman Coulter) to determine total viable cell number. Blood was collected into tubes containing HBSS with 5 mM EDTA, RBC lysed in ACK buffer, washed with media and stained for flow cytometry.

### Flow cytometry

Cells were incubated with αCD16/32 (clone 2.4G2; hybridoma supernatant) and plated on U-bottom 96-well plates at ≤ 3 × 10^6^ cells/well in complete RPMI. Where indicated, cells were stained with Kb-SIINFEKL tetramer APC (NIH tetramer core) at 37°C for 30 min in the presence of αCD8α (53-6.7; Biolegend). For live-dead and surface staining, cells were washed with media, and stained in media for 20 min at RT with Fixable Viability Dye 780 (eBioscience) and surface antibodies for CD19 (6D5, Biolegend), CD8α (53-6.7, Biolegend), CD44 (IM-7; Tonbo), CD127 (A7R34, Tonbo), KLRG1 (2F1/KLRG1, Biolegend), CD45.1 (A20, Biolegend), CD45.2 (104, Biolegend), CD122 (TM-b1, Biolegend). After wash with RPMI, cells were fixed and permeabilized for 45 min at RT in Foxp3 / Transcription Factor 1× Fix/Perm solution (Tonbo), followed by wash with 1× Flow Cytometry Perm Buffer (Tonbo) and intracellular staining for 45 min at RT with intracellular antibodies, including TCF1 (C63D9; Cell Signaling Technology), FOXO1 (C29H4, Cell Signaling Technology), T-bet (4B10, Biolegend), EOMES (Dan11mag; eBioscience). The cells were washed twice and resuspended in 1× Flow Cytometry Perm Buffer (Tonbo). Flow cytometry data was acquired on a four-laser (405, 488, 561, 638 nm) CytoFlex S (Beckman Coulter) and analyzed using FlowJo software (BD Biosciences).

### Bulk RNA sequencing Methods

Congenically marked WT or IL-27R-/- OT1 T cells were flow sorted 3 days after subunit vaccination using a FACSAria Fusion (BD Biosciences) and total RNA was isolated using Qiagen RNeasy columns. RNA was processed for next-generation sequencing (NGS) library construction as developed in the National Jewish Health (NJH) Genomics Facility for analysis with a HiSeq 2500 (Illumina). A SMARTer® Ultra™ Low Input RNA Kit for Sequencing – v4 (Clontech) and Nextera XT (Illumina) kit were used. Briefly, library construction started from isolation of total RNA species, followed by SMARTer 1st strand cDNA synthesis, full length dscDNA amplification by LD-PCR, followed by purification and validation. After that, the samples were taken to the Nextera XT protocol where the sample is simultaneously fragmented and tagged with adapters followed by a limited cycle PCR that adds indexes. Once validated, the libraries were sequenced as barcoded-pooled samples and run on the HiSeq 2500 by the NJH Genomics Facility. FASTQ files were generated using the Illumina bcl2fastq converter (version 2.17).

### Bioinformatic processing of FASTQ files

Nextera adapters were removed from RNA-seq reads with skewer (version 0.2.2) (Jiang et al., 2014). After adapter removal, reads with lengths less than 18 were removed (default setting for skewer). The quality of the reads was assessed using FastQC (version 0.11.5) (Andrews, 2010) before and after read trimming. Reads were mapped with the STAR aligner (version 2.4.1d) (Dobin et al., 2012) to the GRCm38 assembly of the mouse genome using gene annotations from Ensembl version 90 (Ensembl, 2017). Reads mapping to each gene of the Ensembl 90 annotation were counted with the featureCounts program from the Subread software package (version 1.6.2) (Liao et al., 2013). The counts were transformed using the variance-stabilizing transform (VST) in DESeq2 (Love et al., 2014) version 1.20.0. Principal component analysis was done with the scikit-learn library (Pedregosa et al., 2011) version 0.19.2 on the VST of the counts. Hierarchical clustering was done with the clustermap function in Seaborn (Waskom et al., 2014), using the Ward linkage method and Euclidean distances on the VST of the counts. Statistical comparisons were conducted using the likelihood ratio test in the EdgeR package (version 3.24.3) (Robinson et al., 2010) for the R statistical software (version 3.6). The p-values reported were adjusted for testing using the method by Benjamini & Hochberg (Benjamini and Hochberg, 1995).

### Heatmap generation

Normalized counts from DESeq2 were filtered for genes included in the HALLMARK_MYC_TARGETS_V1 dataset from the Molecular Signatures Database (Liberzon et al., 2015). A heatmap visualization of these filtered counts was then generated using the Morpheus software from the Broad Institute.

### Single-cell RNA sequencing analysis (scRNAseq)

#### scRNAseq sample preparation

A small number (500) of congenically marked (CD45.1) OT1 T cells were adoptively transferred into a naïve B6 recipient (CD45.2). These recipients were then immunized intravenously (IV) against ova using our combined adjuvant vaccine (poly I:C, anti-CD40) or infection with LM-ova. T cells were isolated on day 7 post immunization and pooled from 5 mice each prior to cell sorting of CD45.1+ CD8+ cells. Cells from an individual condition (adjuvant or LM) were stained with an oligo-tagged anti-CD45 “hashtag” antibody (Biolegend). About 20,000 cells per were then loaded into Single Cell A chips (10x Genomics) and partitioned into Gel Bead In-Emulsions in a Chromium Controller (10x Genomics). Single-cell RNA libraries were prepared according to the 10x Genomics Chromium Single Cell 3′ Reagent Kits v2 User Guide and sequenced on a NovaSeq 6000 (Illumina).

#### scRNAseq mapping

Reads from scRNA-seq were aligned to mm10 or hashtag oligo sequences and collapsed into unique molecular identifier (UMI) counts using the 10x Genomics Cell Ranger software (version 2.1.0). The sample had appropriate numbers of genes detected (>1000), a high percentage of reads mapped to the genome (>70%), and a sufficient number of cells detected (>1000).

#### scRNA-seq dataset statistical analysis

The Seurat package (version 4.3.0) in R (version 4.0.4) was used for processing, analysis and visualization of the scRNAseq dataset. UMIs with gene counts less than 250, mitochondrial gene content of greater than 15% or expression of multiple hashtag sequences (doublets) were filtered out. Hashtag sequence counts were first normalized using the Seurat function NormalizeData and then demultiplexed using the Seurat function HTODemux which uses K-means clustering. Five mice per group were used for the single cell experiment. Cell read output from Cell Ranger was loaded into Seurat and then Adjuvant or LM-elicited cells were demultiplexed using the level of hashtag sequence. The scRNA-seq dataset was further filtered on the basis of gene numbers, mitochondria gene counts to total counts ratio, and expression of single hashtag sequences. Gene count data was then normalized using Seurat’s NormalizeData function. Top variable genes, principal components analysis (PCA), and uniform manifold approximation and projection for dimension reduction (UMAP) were calculated by the functions: FindVariableGenes, RunPCA, and RunUMAP. Only the top 2000 genes were considered in the PCA calculation and only the top 20 PCs were used in UMAP. Shared nearest-neighbor (SNN) identification and clustering were performed using the functions FindNeighbors and FindClusters using the top 20 PCs with resolution set to 0.3 and k set to 30.

#### cNMF: Identification of Gene Expression Programs

The count matrix was used for conducting non-negative matrix factorization (NMF) through the cNMF method53. This process enabled us to infer both identity and activity programs, along with their respective contributions in each cell. Cells were assigned to the GEP with the highest GEP score and this assignment was added to the metadata of the Seurat object. To determine the genes associated with each program, we plotted the gene ranks (ranging from most associated to least associated) against the gene_spectra_score output from the cNMF analysis. This ranked gene expression was utilized in GSEA using the fGSEA (version 1.24.0) package in R. Both positive and negative normalized enrichment scores (NES) were obtained and scaled NES were graphed for all gene sets with an adjusted p value > 0.05.

#### Gene regulatory network Interference

To deduce gene regulatory networks, we employed pySCENIC from a pre-built singularity container, aertslab/pyscenic:0.12.1, a tool utilizing cis-regulatory motif analysis to identify potential transcription factors (TFs) that might govern a cluster of co-expressed genes within individual cells52. pySCENIC was run using the –mask-dropouts flag and a normalized enrichment score threshold of 2 to help mitigate the effects of the varying degrees of sparsity across the data sets we generated. The initial step involved generating modules composed of transcription factors and co-expressed genes using GRNboost2 (Ref 107). These modules were pruned to remove indirect targets that lacked significant enrichment for the corresponding TF motif within ±10 Kb from the transcription starting site of the putative target (cisTarget). This process yielded a collection of transcription factor regulons. Considering the inherent stochasticity in gene regulatory network inference using GRNBoost2, each run of pySCENIC may yield different quantities of regulons, along with distinct target genes associated with each TF. To mitigate this variability, we performed 100 pySCENIC runs and retained regulons present in 100% of the runs. We also removed regulons that did not have at least 5 target genes defining the regulon activity. Due to the high degree of noise in target genes, we retained target genes that appeared within a regulon in at least 95% of the runs. Furthermore, each target gene also had to overlap with the union of all possible retained ranked gene expression targets across all GEPs generated from cNMF. To identify regulons that were specific to the underlying biology of our cell types and GEPs, we calculated the AUC scores using the R package AUCell, located in the pySCENIC container, for each regulon based on the pruned target gene list. A regulon was deemed specific to a defined cell population if at least 20% of the cells within the annotated population scored in the 90th percentile of the overall AUC score for all cells.

#### scRNA analysis packages used

*The R packages Seurat (version 4.3.0), dplyr (version 1.1.3), fgsea (1.24.0), tidyverse (2.0.0), DESeq2 (1.38.3), SCENIC (1.3.1), SCpubR (version 2.0.2), ggplot (version 3.4.4), and pheatmap (version 1.0.12) were used for the analysis and graphing of scRNA data*.

#### Statistical Analysis

GraphPad Prism (version 9.3.0, GraphPad) was used for all statistical analyses. Figure legends detail the number of experimental replicates and n-values. Unless noted, data presented are means ± SEM. Significance was defined using unpaired Student’s t-test with Welch’s correction or analysis of variance (ANOVA). Significance was denoted as follows: ns, not significant (p > 0.05) or significant with a p-value less than 0.05.

---

**Figure S1.** Cluster-defining genes. Heatmap of top 5 expressed genes expressed as a factor of time x challenge (A) or UMAP cluster (B).

**Figure S2.** Determining genes associated with cNMF derived Gene Expression Programs (GEPs). Gene ranks (sorted most to least associated, x-axis) are displayed against their gene_spectra_score output from the cNMF analysis (y-axis) as black dots. The slope at the first elbow point in the fitted sigmoid curve (red line) was calculated as the minimum threshold for genes to be retained in a given GEP. The same slope (grey dashed line) was applied to every GEP to prevent bias in ranked gene selection, as the gene ranking between GEPs are not comparable and relative to each GEP.

**Figure S3.** GSEA of Gene Expression Profiles. GSEA from GEP-defining genes for GEPs 1-3 against the mouse C7 Immunologic gene signature gene sets. All CD8+ T cell associated gene sets in C7 identified as statistically significant are shown, normalized by row.

**Figure S4.** GO pathway and SCENIC analysis of GEPs 1-3. A. GO pathway analysis from of GEP-defining genes, grouped by cellular function. See Table S2 for GO pathways in heatmap. B. Per cell SCENIC-derived regulon scores, ordered by assigned GEP. See Table S3 ordered list of heatmap regulons C. Mean SCENIC regulon scores across all cells/GEP form A, ordered by assigned GEP, normalized by row.

**Figure S5.** WT vs IL-27R KO T cell responses to protein and mRNA vaccination. A and B. CD98 and CD71 gMFI of WT and IL-27R-/- T cells 5 days after vaccination or LM challenge for the experiment shown in Figure 4I and J. C. WT CD45.1/2+, and IL-27R-/- CD45.1/1, OT1 T cells were co-transferred into CD45.2/2 B6 recipients which were then immunized with mRNA-LNPs, the mRNA encoding for ova expression. Blood was analyzed at the time points shown for the representation of WT vs IL-27R-/- out of total transferred T cells.

**Table S1.** Ranked GEP-defining genes after fitting to a sigmoid curve to calculate the minimum threshold for genes to be retained in a given GEP.

**Table S2.** Ordered list of gene sets shown in Fig. 2L.

**Table S3.** Ordered list of GO-BP’s as shown in Fig. S4.

**Table S4.** Mean regulon scores of all statistically significant regulons from SCENIC analysis shown in Fig S4.

