## Supplementary figures and images for "IL-27 Stablizes Myc-Mediated Transcription In Memory-Fated, Vaccine-Elicited CD8+ T Cells"

### supplementary Figure 1

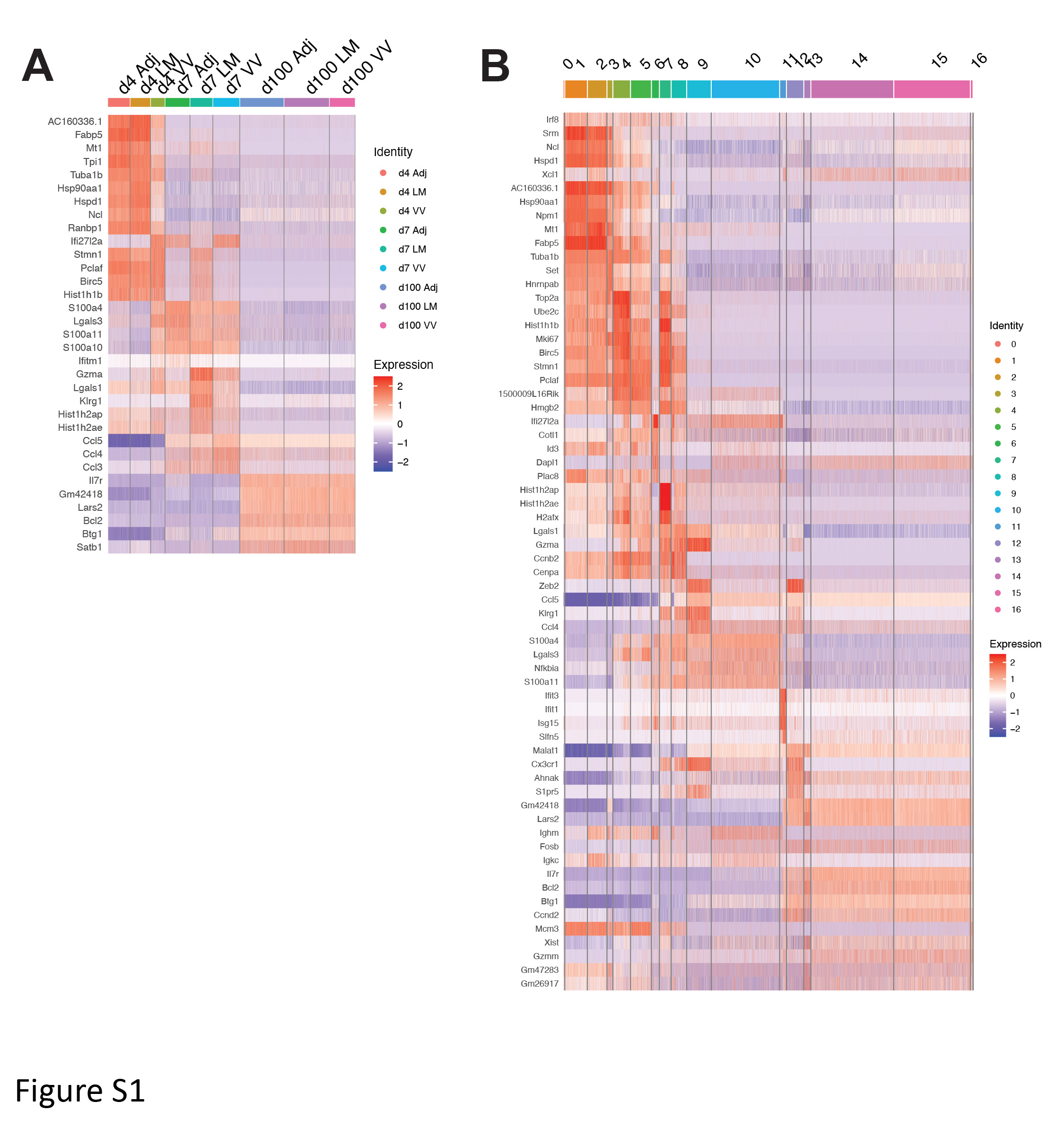

### supplementary Figure 2

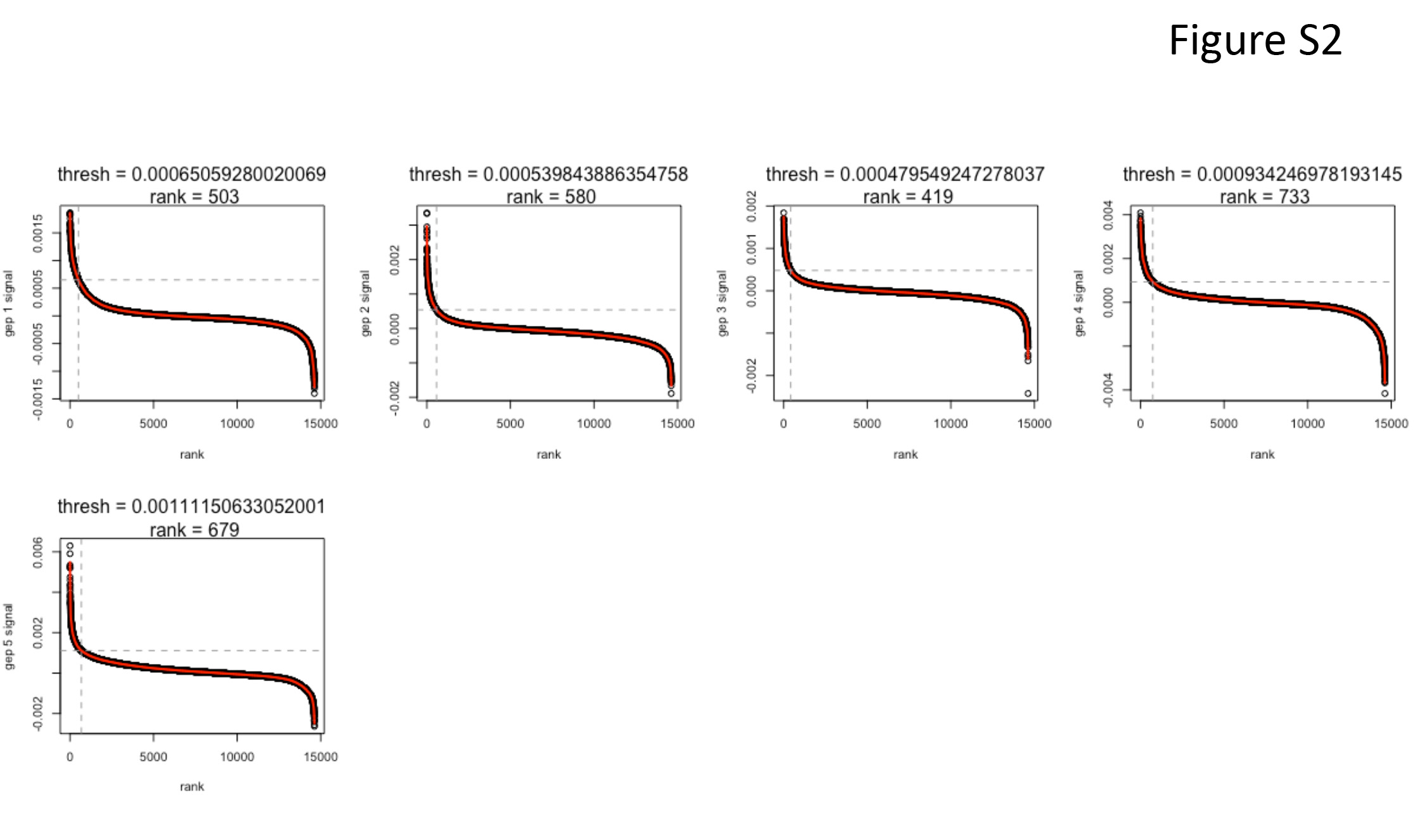

### supplementary Figure 3

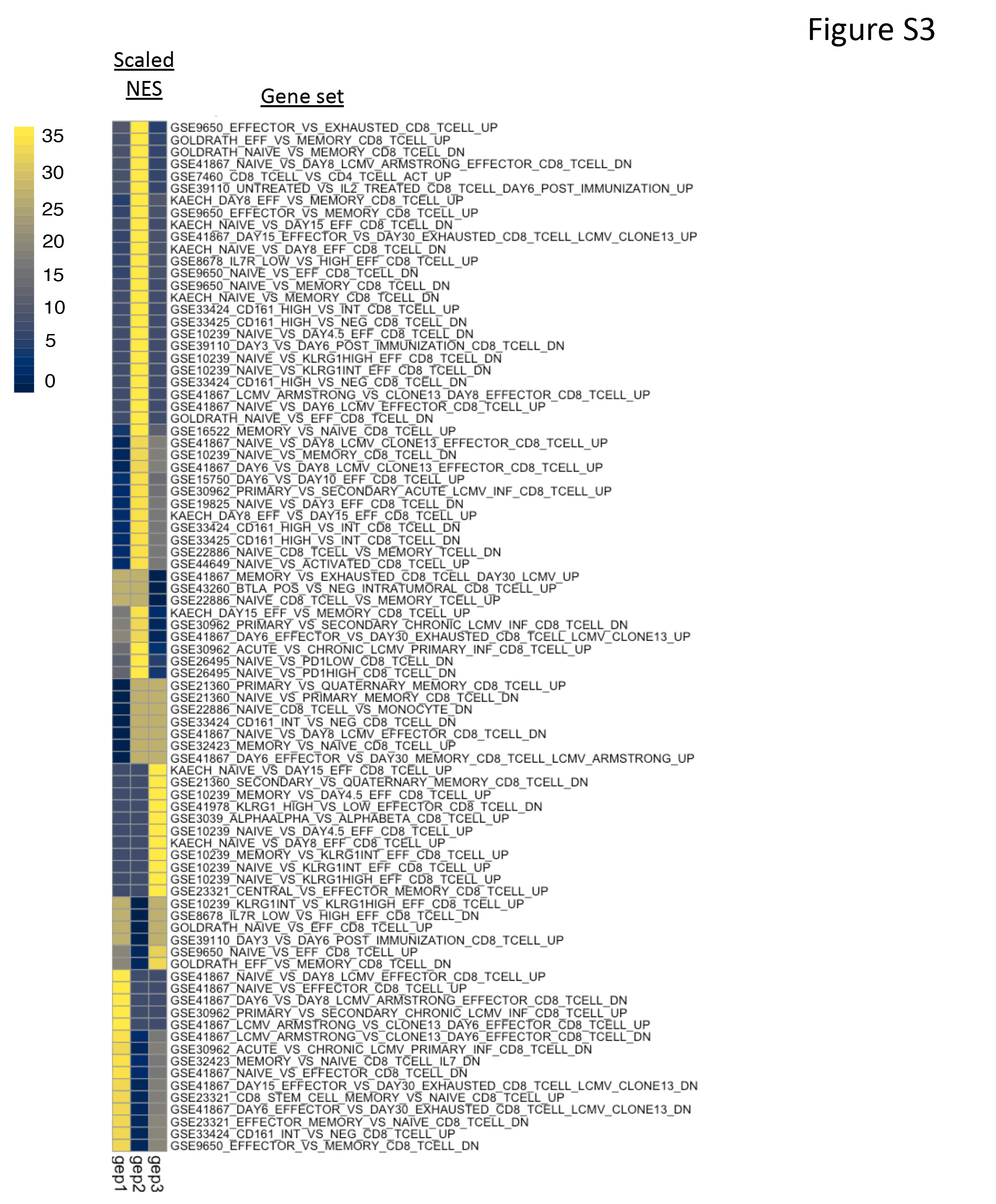

### supplementary Figure 4

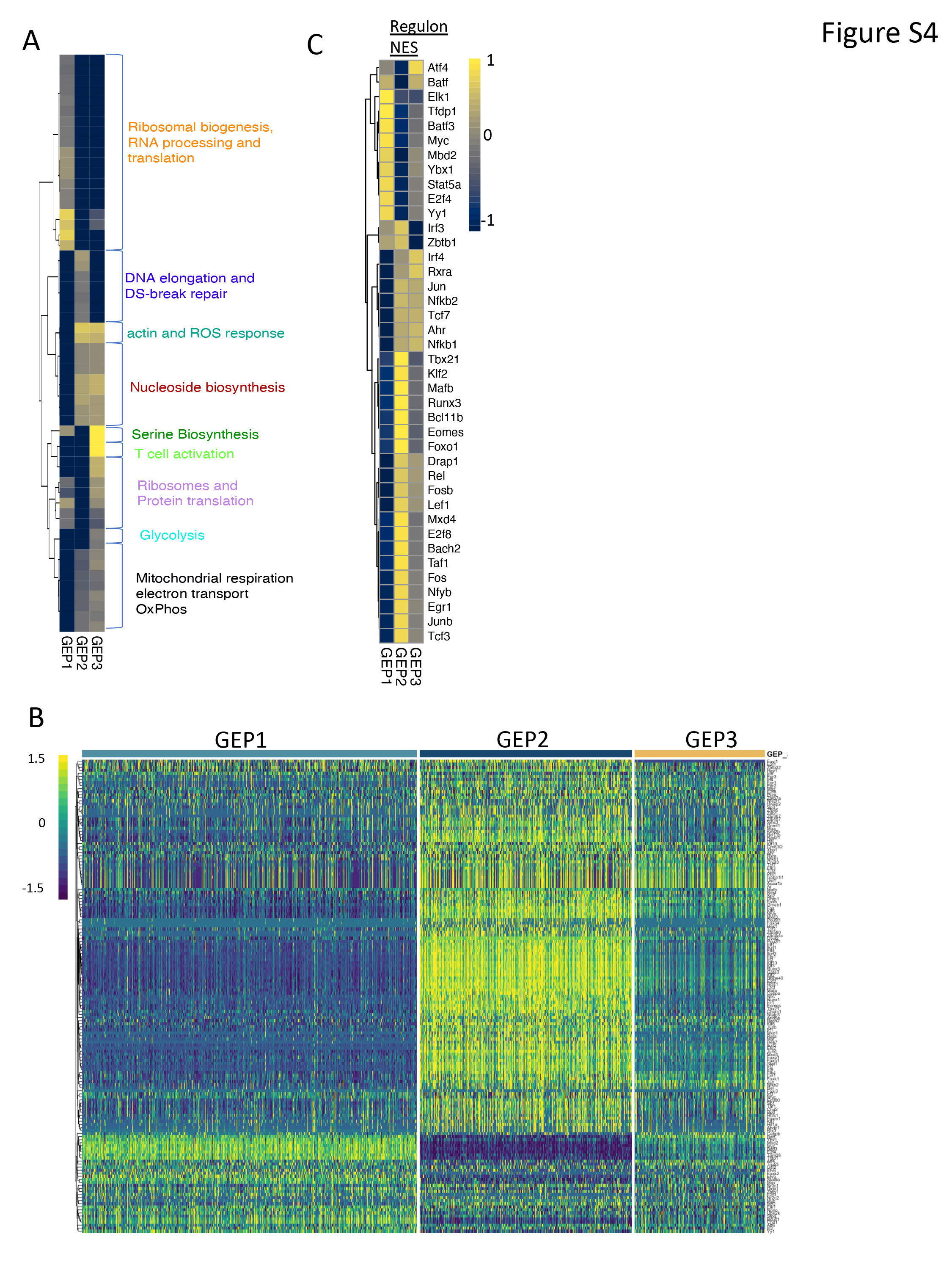

### supplementary Figure 5

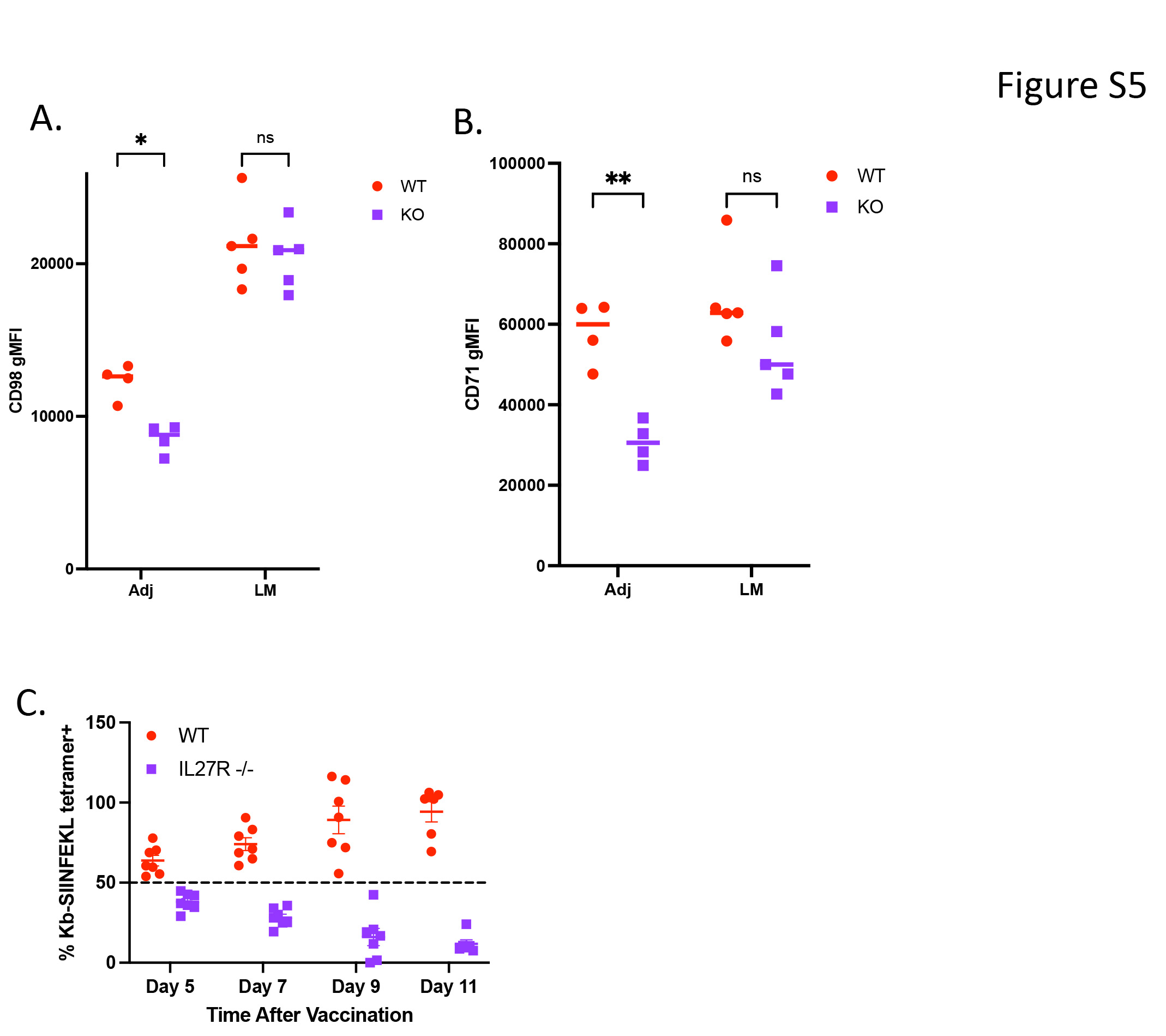
